# Ontogenic, phenotypic, and functional characterization of XCR1^+^ dendritic cells leads to a consistent classification of intestinal dendritic cells based on the expression of XCR1 and SIRPα

**DOI:** 10.1101/004648

**Authors:** Martina Becker, Steffen Güttler, Annabell Bachem, Evelyn Hartung, Ahmed Mora, Anika Jäkel, Andreas Hutloff, Volker Henn, Hans W. Mages, Stephanie Gurka, Richard A. Kroczek

## Abstract

In the past, lack of lineage markers confounded the classification of dendritic cells (DC) in the intestine and impeded a full understanding of their location and function. We have recently shown that the chemokine receptor XCR1 is a lineage marker for cross-presenting DC in the spleen. Now we provide evidence that intestinal XCR1^+^ DC largely, but not fully, overlap with CD103^+^ CD11b^-^ DC, the hypothesized correlate of “cross-presenting DC” in the intestine, and are selectively dependent in their development on the transcription factor Batf3. XCR1^+^ DC are located in the villi and epithelial crypts of the lamina propria of the small intestine, the T cell zones of Peyer’s Patches, and in the T cell zones and sinuses of the draining mesenteric lymph node. Functionally, we could demonstrate for the first time that XCR1^+^ / CD103^+^ CD11b^-^ DC excel in the cross-presentation of orally applied antigen. Together, our data show that XCR1 is a lineage marker for cross-presenting DC also in the intestinal immune system. Further, extensive phenotypic analyses reveal that expression of the integrin SIRPα consistently demarcates the XCR1^-^ DC population. We propose a simplified and consistent classification system for intestinal DC based on the expression of XCR1 and SIRPα.

## Introduction

The intestinal immune system has to discriminate between harmless food proteins, commensal bacteria colonizing the gut, and dangerous pathogens. Dendritic cells play a central role in orchestrating the appropriate immune responses. Conventional dendritic cells (DC) reside in the lamina propria (LP) of the small and large intestine, in the scattered lymphoid follicles and Peyer’s patches (PP), and in lymph nodes draining the intestine, such as the mesenteric lymph nodes (MLN). In the past, any subdivision of intestinal DC into functional subpopulations was controversial because DC-specific markers were lacking and other surface molecules used for classification were found to be regulated or were also present on macrophages (Milling et al., 2010; Pabst and Bernhardt, 2010; Rescigno, 2010; Hashimoto et al., 2011; Mowat and Bain, 2011; Bogunovic et al., 2012). A major step forward was the combined use of antibodies directed to the integrins CD103 and CD11b, which allowed to define four DC subsets, CD103^+^ CD11b^-^, CD103^+^ CD11b^+^, CD103^-^ CD11b^+^, and CD103^-^ CD11b^-^ (Annacker et al., 2005; Johansson-Lindbom et al., 2005; Schulz et al., 2009), which in the MLN were further grouped into resident and migratory DC (Ohl et al., 2004). Because of open questions regarding the subdivision of DC in the intestine, only very few studies on antigen presentation were performed with intestinal DC.

Most of the work on the function of DC has been performed with splenic DC populations. These studies demonstrated that all DC can present exogenous antigen to CD4^+^ T cells (classical presentation), while only a subset excels in the cross-presentation of antigen to CD8^+^ T cells (den Haan et al., 2000; Pooley et al., 2001). Cross-presentation is a central element in the activation of CD8^+^ T cells to cytotoxic effector cells, and thus of major importance in the defense of certain infections and in the elimination of tumors (Bevan, 2006; Shen and Rock, 2006; Villadangos et al., 2007). Based on the commonly used classification of splenic DC, numerous studies established that CD8α^+^ DC excel over CD4^+^ DC and double negative DC in their capacity to cross-present (cell-associated) antigen. Because of the different DC classification systems used it remained unclear whether these splenic CD8α^+^ DC have a correlate among the intestinal DC.

A major advance for a unified classification of DC throughout the immune system was brought by analyses on role of transcription factors (TF) in the differentiation of DC. These studies revealed that CD8α^+^ DC in the spleen and CD103^+^ CD11b^-^ DC in the intestine and other organs were specifically absent in animals deficient for the TF Batf3 (Hildner et al., 2008; Edelson et al., 2010). These findings strongly indicated that CD8α^+^ DC in the spleen and the lymphoid organs correspond to CD103^+^ CD11b^-^ DC in tissues, and together represent a separate DC lineage of cross-presenting DC. The intestinal CD103^+^ CD11b^-^ DC were thus subsequently also termed “CD8α-like DC” or “Batf3-dependent DC”.

Recently, we and others have recognized that the chemokine receptor XCR1 is exclusively expressed on murine (Dorner et al., 2009) and human DC (Bachem et al., 2010; Crozat et al., 2010). In the murine system, expression of XCR1^+^ DC was found to be restricted to CD8α^+^ DC in the spleen and CD8α-like DC in peripheral organs (Dorner et al., 2009; Crozat et al., 2011; Bachem et al., 2012). However, in a subsequent more detailed analysis it became apparent that splenic CD8α^+^ DC are not identical to XCR1^+^ DC, since these two populations only overlap (Bachem et al., 2012). We could firmly establish that splenic DC expressing both XCR1 and CD8α+, or only XCR1, belong to the Batf3-dependent, antigen-cross-presenting DC (Bachem et al., 2012). In contrast, CD8α^+^ DC lacking XCR1 on the cell surface are a clearly different population, which i) is independent of Batf3, ii) has a distinct gene expression program (Bar-On et al., 2010), and iii) is functionally different, as it is unable to cross-present antigen (Bachem et al., 2012). With this work it became clear that the identified lineage of Batf3-dependent DC in the spleen (and possibly other organs) is truly represented by DC expressing XCR1. At the same time, this work determined that “CD8α^+^ DC” are in reality a phenotypically and functionally heterogeneous population.

With the present study we analyzed the development, phenotype, localization, and function of XCR1^+^ cells in the intestinal immune system to find out whether they correspond to XCR1^+^ DC in the spleen. Although our work was focused on XCR1^+^ DC, most of the experiments also yielded information on XCR1^-^ DC, which were used for comparison. The results demonstrate that the expression of XCR1 also in the intestinal immune system consistently demarcates the lineage of Batf3-dependent, antigen cross-presenting DC. At the same time, we show that intestinal XCR1^+^ DC strongly overlap, but are not fully congruent with CD103^+^ CD11b^-^ DC.

Further, our extensive phenotypic studies demonstrate that XCR1 and SIRPα delineate two mutually exclusive DC populations, which together encompass essentially all conventional DC in the intestine. Based on these results we propose a new and simplified classification system for conventional DC in the intestine based on only two surface markers, XCR1 and SIRPα.

## Materials and methods

### Mice and Flt3 ligand treatment

Unless indicated otherwise, 8-10 week-old C57BL/6 female mice were used for cell isolation and immunohistological analyses. CX_3_CR1^GFP^ (Jung et al., 2000), Langerin^EGFP^ mice (Kissenpfennig et al., 2005), B6.XCR1-lacZ^+/+^ (The Jackson Laboratories) and Batf3-deficient mice (Hildner et al., 2008) were on the C57BL/6 background. OT-I TCR-transgenic mice were crossed onto the B6.PL background to allow identification of CD8^+^ T cells using the CD90.1 marker. For Flt3 ligand treatment, C57BL/6 mice were injected with 1×10^6^ B16 cells secreting Flt3 ligand (Mach et al., 2000) in 100 µl PBS s.c. All mice were bred under specific pathogen-free conditions in the animal facility of the Federal Institute for Risk Assessment (Berlin, Germany). All animal experiments were performed according to state guidelines and approved by the local animal welfare committee.

### Antibodies

Hybridomas producing mAb recognizing CD4 (clone YTS 191.1), CD8α (53-6.72), CD11b (5C6), CD11c (N418), CD16/32 (2.4G2), CD45R/B220 (RA3-6B2), DCIR2 (33D1), MHCII (M5/114.15.2), and NK1.1 (PK136) were obtained from ATCC, CD90.1 (OX-7) from ECACC. MAb to CD103 (M290), CD172a/ SIRPα (P84), and CD11c (HL3) were from BD Biosciences, to CD69 (H1.2F3), CD45 (30F11) and CCR7 (4B12) from eBioscience, and to CD3 (17A2), F4/80 (BM8) and CD45R/B220 (RA3-6B2) from BioLegend. Anti-XCR1 (MARX10 (Bachem et al., 2012)) and anti-Clec9A/DNGR-1 (clone 24/04-10B4 (Caminschi et al., 2008)) antibodies were used. Anti-CD3 (KT3) was generously provided by H. Savelkoul, anti-CD25 (2E4) by E. Shevach, and anti-DEC 205 (NLDC-145, CD205) by G. Kraal.

### Cell isolation

For isolation of small intestinal LP DC, the small intestine was freed from fat and PP, opened longitudinally, and stirred in PBS, 2% FCS, 1 mM EDTA, 1 mM DTT for 7 min at 37°C. After additional stirring under the same conditions without DTT, epithelial cells in solution were discarded, intestinal tissue was minced, and stirred in 500 µg/ml Collagenase VIII (in some experiments Collagenase D was used instead in an attempt to improve staining of Clec9a, both from Sigma) and 20 µg/ml DNAse I (Roche) for 30 min at 37°C; thereafter cells were mashed through a 70 µm nylon sieve (BD Falcon). For isolation of DC from lymphoid tissues, MLN and PP were ruptured and digested with Collagenase D (500 µg/ml) and DNase I (20 µg/ml, both Roche) for 15 min at 37°C in RPMI 1640 containing 2% FCS (low endotoxin, Biochrom); EDTA (10 mM) was added for additional 5 min and cells were mashed through a 70 µm nylon sieve. For staining of DC from LP and PP, low density cells from these tissues were enriched by centrifugation over a 1.073 g/ml density gradient (Nyco-Prep, Axis-Shield). For flow sorting of DC from LP and MLN, low density cells were enriched and magnetically sorted with CD11c microbeads (Miltenyi Biotec). Splenocytes were obtained by mashing spleens through 70 µm cell sieves into PBS, followed by erythrocyte lysis with ACK Buffer (155 mM NH_4_Cl, 10 mM KHCO_3_, 0.1 mM EDTA).

### Flow cytometry and flow sorting

Antibodies were titrated for optimal signal-to-noise ratio. To block unspecific binding to Fc-receptors, cells were pre-incubated with 100 µg/ml 2.4G2 mAb for flow cytometry and in addition with 50 µg/ml purified rat Ig (Nordic) for flow sorting. Doublets and autofluorescent cells were excluded from the analysis. In all organs, DC were identified as CD11c^+^ MHCII^+^ Lin^-^ F4/80^-^ cells, in LP and PP DC were additionally defined using CD45; the lineage cocktail contained mAb directed to CD3 and B220. Standard staining with mAb was in PBS, 0.25% BSA, 0.1% NaN_3_ for 20 min on ice, staining for Clec9A was in the same buffer for 20 min at 37°C. For exclusion of dead cells, 4‘,6-diamidino-2-phenylindole (DAPI) was added 5 min before measurement. Data were acquired on a LSRFortessa flow cytometer (BD Biosciences) and analyzed using FlowJo (Tree Star Inc.). For analysis of surface receptor expression gates were set according to the appropriate isotype/FMO controls. Flow sorting of small intestinal DC (CD11c^+^ MHCII^+^ CD45^+^ Lin^-^ F4/80^-^), migratory MLN DC (CD11c^+^ MHCII^high^ Lin^-^ F4/80^-^), and resident MLN DC (CD11c^+^ MHCII^low^ Lin^-^ F4/80^-^) was based on their expression of CD103 and XCR1 and performed on a FACSAriaII (BD Biosciences).

### Histology

For histological β-galactosidase analysis, tissues from homozygous B6.XCR1-lacZ^+/+^ mice were fixed with 0.1% glutaraldehyde plus 4% paraformaldehyde in PBS for 4 h at RT, immersed in 10% sucrose overnight, and snap-frozen in 0.9% NaCl. Cryostat sections (15 µm) were washed with cold PBS (pH 7.4) for 5 min after thawing, incubated with X-Gal staining solution (Sanes et al., 1986) overnight at 37°C, washed with PBS, and counterstained with Neutral Red.

### Cross-presentation assay

C57BL/6 mice were fed with 25 mg ovalbumin (OVA, Sigma-Aldrich) in 500 µl PBS by gavage. 17 hours after oral application of OVA, DC subsets were flow sorted to high purity (> 98.5%). OT I CD8^+^ T cells were enriched from spleens of OT I mice by magnetically depleting cells expressing CD4, CD11b, CD11c, B220, or NK1.1 (Miltenyi Biotec). Preparations of resting OT I T cells (confirmed by negativity for CD25 and CD69) were labeled with CFSE (Molecular Probes, 5 µM, 15 min, 37°C). For cross-presentation assays, 1×10^5^ CFSE-labeled OT-I T cells were co-cultured with titrated numbers (3,750-30,000) of DC subsets in 200 µl RPMI medium containing 10% FCS, 50 µM 2 mercaptoethanol, 1 mM sodium pyruvate, non-essential amino acids, and 100 µg/ml penicillin/streptomycin in 96-well round-bottomed plates (Nunc) for 2.5 days. Thereafter, proliferation of OT I T cells was determined in the CFSE dilution assay after gating on CD90.1 cells. For positive control, sorted DC were incubated with 1 µM of the OVA peptide SIINFEKL, and co-cultured with CFSE-labeled OT I T cells for 2.5 days.

## Results

### Phenotype of XCR1^+^ DC in the lamina propria, Peyers’ patches, and mesenteric lymph nodes

To determine the phenotype of XCR1^+^ DC in the intestinal immune system at steady state, mononuclear cells were obtained from the lamina propria (LP) of the small intestine, Peyer’s patches (PP), and mesenteric lymph nodes (MLN). Conventional DC, defined as lineage-negative, F4/80-negative, CD11c^hi^ MHCII^hi^ cells by flow cytometry, were then gated into four populations based on the expression of the integrins CD103 (Itgae) and CD11b. In the LP, essentially all CD103^+^ CD11b^-^ DC expressed XCR1, while almost all CD103^+^ CD11b^+^ and CD103^-^ CD11b^+^ DC were negative (Fig. 1). Interestingly, a fraction (around 10%) of the small population of CD103^-^ CD11b^-^ (double-negative) DC found in the small intestine also expressed XCR1 (Fig. 1). In the PP, a large majority (around 80%) of CD103^+^ CD11b^-^ DC expressed XCR1, while all other populations resembled in their XCR1 expression profile LP cells (Fig. 1). In MLN, expression levels of MHCII were used to further discriminate resident from migratory DC (Ohl et al., 2004). Migratory DC subpopulations closely resembled in their XCR1 expression pattern LP DC (Fig. 1). Different from LP, PP, and migratory MLN DC, the CD103^-^ population in the resident MLN DC contained a substantial proportion of XCR1^+^ DC (Fig. 1). Taken together, our results, based on the currently popular subdivision of intestinal DC using CD103 and CD11b as markers, demonstrated a large but clearly not full overlap between XCR1^+^ DC and CD103^+^ CD11b^-^ DC.

**Figure 1.**
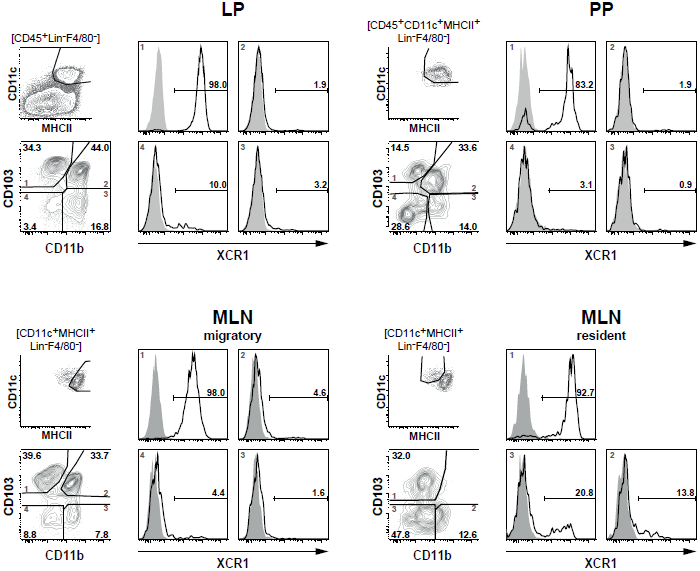
Expression of XCR1 on DC in the lamina propria, Peyer’s patches, and mesenteric lymph nodes. DC from the LP, PP, and MLN of C57BL/6 mice were enriched by digestion and density gradient centrifugation of the tissues, stained for CD11b and CD103, and counter stained with XCR1-specific mAb MARX10. DC from MLN were separated into resident and migratory DC based on their MHCII expression levels. For analysis, the gates were set on live CD45^+^ Lin^-^ F4/80^-^ CD11c^+^ MHCII^+^ cells. Expression of XCR1 is shown on CD103^+^ CD11b^-^ (left upper quadrants), CD103^+^ CD11b^+^ (right upper quadrants), CD103^-^ CD11b^+^ (right lower quadrants) and CD103^-^ CD11b^-^ (left lower quadrants) DC. The background staining was determined with homozygous B6.XCR1-lacZ^+/+^ mice lacking XCR1 (gray). The results shown are representative of three experiments with three animals each.

To further define the phenotype of XCR1^+^ DC at the examined locations, expression of XCR1 was correlated to a greater number of surface molecules known to be expressed on DC in the intestine. In the steady state, expression of XCR1 was found to be highly correlated with CD8α on DC in the LP, PP, and MLN (Fig. 2). Only a small population of resident MLN DC expressed CD8α, but was negative for XCR1 (Fig. 2), as earlier found in the spleen (Bachem et al., 2012). When detectable, expression of Clec9A/DNGR-1 on DC was highly correlated with XCR1 in all anatomical sites. Surface presence of XCR1 corresponded well with CD205 in the LP and MLN, but less so in the PP. XCR1^+^ DC usually co-expressed the integrin CD103, which in resident MLN DC was found to be partly downregulated. XCR1^+^ DC were low/ negative for CX_3_CR1/ fractalkine receptor, and with the exception of a population of resident MLN DC, also negative for CD207/ langerin. In all instances, XCR1^+^ DC were negative for 33D1/ DCIR2 and CD11b, but neither of these molecules was anti-correlated. Finally and importantly, only CD172a/SIRPα was clearly anti-correlated with XCR1 and also encompassed all XCR1^-^ DC.

**Figure 2.**
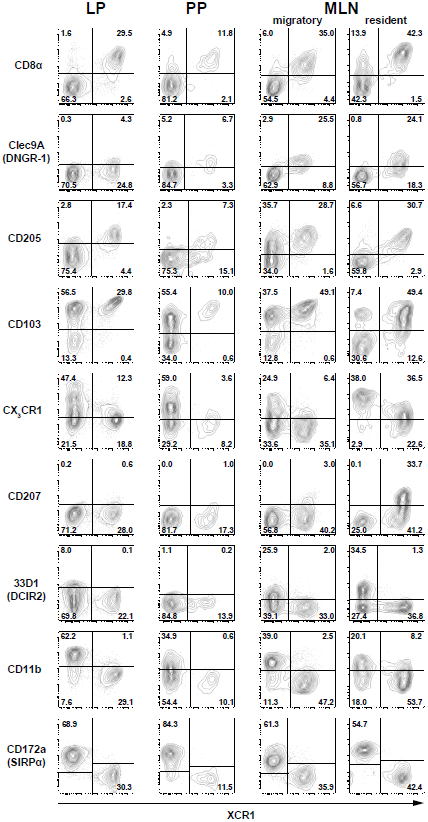
Correlation of XCR1 expression with different DC surface molecules. DC from the LP, PP, and MLN of C57BL/6 wt mice, and heterozygous CX_3_CR1^GFP^ or Langerin^EGFP^ (CD207) mice were enriched by digestion and density gradient centrifugation and double-stained for detection of XCR1 and the indicated surface molecules. For analysis, the gates were set on live CD45^+^ Lin^-^ F4/80^-^ CD11c^+^ MHCII^+^ cells. The results shown are representative of three experiments with three animals each.

In summary, none of the surface molecules examined was fully correlated with XCR1 on DC in all anatomical locations. The best overall correlation of XCR1 was seen with CD8α and with Clec9A/DNGR-1 (when detectable), indicating a functional link between these three receptors. Further, XCR1 was substantially, but not fully, correlated with CD205 in all tissues. On the other hand, a perfect anti-correlation could be observed between XCR1 and SIRPα in all anatomical sites, which was not the case between XCR1 and CD11b.

### Expansion of intestinal DC populations under the influence of Flt3 ligand

Flt3 ligand is a growth factor described to play a key role in the physiological expansion of classical CD8^+^ DC (Maraskovsky et al., 1996). We have previously observed that Flt3 ligand expanded XCR1^+^ DC in the spleen around 20-fold, while XCR1^-^ DC were expanded only around 4-fold (Bachem et al., 2012). In order to determine whether the same phenomenon can be observed in the intestinal immune system, mice were exposed to Flt3 ligand and the composition of the various DC populations was determined by flow cytometry. Under the influence of Flt3 ligand, both XCR1^+^ CD103^+^ and XCR1^-^ CD103^-^ increased in frequency, and this was accompanied by a substantial relative reduction of XCR1^-^ CD103^+^ DC in all anatomical sites (Fig. 3, flow cytometry histograms). In terms of absolute cell numbers, all DC populations increased under the influence of Flt3 ligand, with biggest changes in PP and MLN. There, XCR1^+^ CD103^+^ and XCR1^-^ CD103^-^ DC expanded both around 30-fold and thus more than XCR1^-^ CD103^+^ DC (10-fold), while the most dramatic change was seen with the usually minute population of XCR1^+^ CD103^-^ DC (up to 400-fold increase). Thus, the effects of Flt3 ligand on the expansion of intestinal DC subsets were more complex than in the spleen, with a massive expansion of XCR1^+^ CD103^-^ DC, but a similar expansion of XCR1^+^ CD103^+^ and XCR1^-^ CD103^-^ DC.

**Figure 3.**
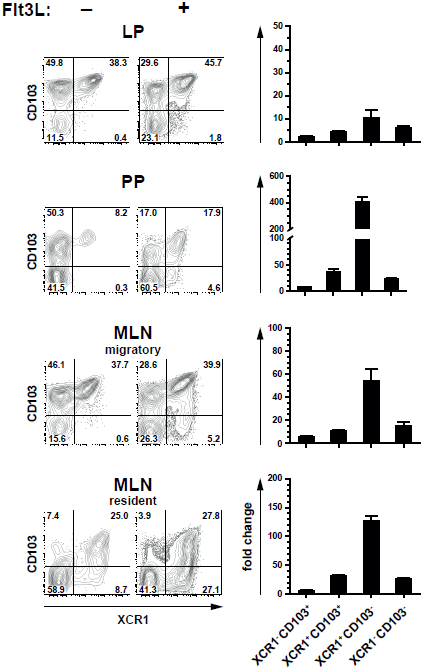
Expansion of intestinal XCR1^+^ DCs by the growth factor Flt3 ligand. C57BL/6 mice were exposed to Flt3 ligand for 9 days in vivo. Thereafter, DC from the LP, PP, and MLN (CD45^+^ Lin^-^ F4/80^-^ CD11c^+^ MHCII^+^ cells) were analyzed for expression of CD103 and XCR1, and compared to unexposed controls (flow cytometry histograms). The bar graphs represent the fold increase in total numbers of the indicated DC subsets in relation to unexposed controls. The data shown are representative of two independent experiments (mean ± SEM; n = 3).

### XCR1^+^ DC in the intestinal immune system are Batf3-depen**dent**

The development of CD8α^+^ DC has originally been described to be dependent on the TF Batf3 (Hildner et al., 2008). However, we recently found that in the spleen only CD8^+^ XCR1^+^ DC were dependent on this TF, but not the CD8α^+^ DC population negative for XCR1, which has a clearly different gene expression profile and function (Bar-On et al., 2010; Bachem et al., 2012). In order to determine the influence of Batf3 on the development of intestinal DC, we analyzed Batf3-deficient animals on the C57BL/6 background. In all anatomical sites Batf3-deficiency essentially resulted in the absence of XCR1^+^ DC, clearly demonstrating that Batf3 is required for the development of XCR1^+^ DC in the gut (Fig. 4).

**Figure 4.**
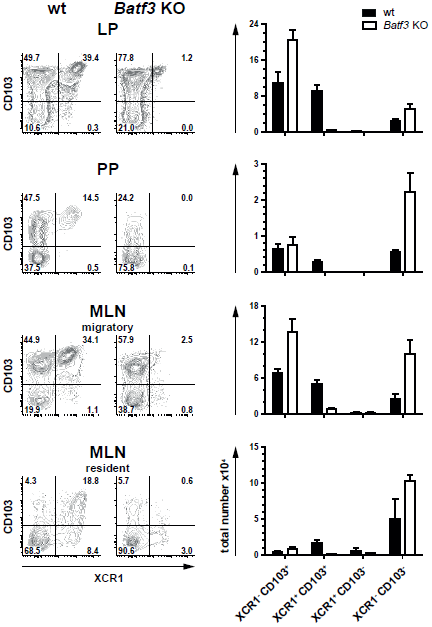
Development of intestinal XCR1^+^ DCs is dependent on the transcription factor Batf3. DC (CD45^+^ Lin^-^ F4/80^-^ CD11c^+^ MHCII^+^) from LP, PP, and MLN cells of C57BL/6 wt controls and Batf3-deficient mice were stained for XCR1 and CD103 (left part of figure). Total numbers of the indicated DC subsets obtained from wt (black bars) and Batf3-deficient mice (open bars) are shown (right part of figure). The results shown are representative of two experiments (mean ± SEM; n = 3).

### Positioning of XCR1^+^ DC in the lamina propria, Peyer’s patches, and mesenteric lymph nodes

Due to the absence of specific markers, an unequivocal localization of DC subsets in lymphoid tissues or organs was very challenging in the past. To overcome these difficulties, we attempted to use the XCR1-specific mAb MARX10 for histological analyses of gut tissues, but this approach gave a high background preventing a clear discrimination from false positive signals. In the next approach, we used B6.XCR1-lacZ^+/+^ mice, in which the XCR1 gene is replaced by the *LacZ* reporter gene, to localize XCR1^+^ DC. Histochemical analysis of β-galactosidase activity in the small intestine gave signals in the lamina propria of the villi (Fig. 5a). In addition, multiple signals were consistently detectable in the epithelial crypts (Fig. 5a). In PP, cells with β-galactosidase activity could be found in the T cell zones, with some clustering in the interfollicular region (Fig. 5b). Interestingly, no signals for XCR1^+^ were detectable in the subepithelial dome of the PP, where CD11c^+^ cells are also known to localize (Kelsall and Strober, 1996). In the MLN, XCR1 signals were seen in the T cell zones and apparently in sinuses (Fig. 5c), similar to the results obtained with axillary LN earlier (Dorner et al., 2009). All of these signals were not present in staining controls (Fig. 5d), indicating that they truly represent cells expressing XCR1 mRNA.

**Figure 5.**
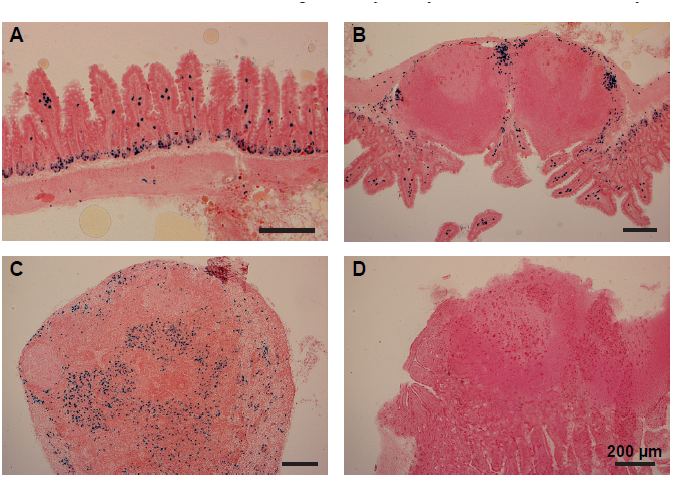
Positioning of XCR1-expressing cells in the lamina propria, Peyer’s patches, and mesenteric lymph nodes. Distribution of XCR1^+^ cells was determined in tissues of homozygous B6.XCR1-lacZ^+/+^ reporter mice using X-gal, a chromogenic substrate for β-galactosidase. (A) Lamina propria of the small intestine, (B) Peyer’s patch, (C) mesenteric lymph node. (D) Represents a staining control using intestinal tissue from wt C57BL/6 mice containing a Peyer’s patch and adjacent portions of the small intestine.

### No apparent involvement of the chemokine receptor XCR1 in the migration of DC to mesenteric lymph nodes

In steady state, intestinal DC constantly migrate from the gut to the MLN in a CCR7-dependent fashion (Jang et al., 2006; Worbs et al., 2006), and this migration is further increased under inflammatory conditions (MacPherson et al., 1995; Turnbull et al., 2005; Schulz et al., 2009). Since XCR1 is also a chemokine receptor, we sought to determine any involvement of XCR1 in the migration of DC from the gut to the MLN, where the ligand XCL1 is secreted by NK cells at steady state and at high levels by activated CD8^+^ T cells, NK cells, and NKT cells (Dorner et al., 2002; Dorner et al., 2004) (own unpublished data). In the first step, expression of XCR1 was correlated with CCR7 under various conditions. At steady state, CCR7 could not be detected on DC in the LP, PP, or resident DC in the MLN, but could be found on the majority of migratory DC (Fig. 6). After intraperitoneal (i.p.) administration of LPS, CCR7 became detectable on around 70% of DC in the PP, but not on LP DC. Under inflammatory conditions, CCR7 was also present on 50-60% of MLN DC, which, due to their uniformly increased levels of MHCII, no longer could be subdivided into resident or migratory DC. In all instances in which CCR7 was detected, essentially all XCR1^+^ DC co-expressed CCR7, suggesting that XCR1 could be involved in the migration of XCR1^+^ DC into the MLN. We therefore performed a series of analyses comparing wt mice with mice deficient for XCL1 (Dorner et al., 2009), the unique chemokine ligand of XCR1. In these experiments, absence of XCL1 did not change the relative representation of the various DC populations in the MLN at steady state or under inflammatory conditions, or their CCR7 expression (own unpublished data). These functional experiments largely excluded an involvement of the XCL1-XCR1 axis in the immigration of DC into the MLN.

**Figure 6.**
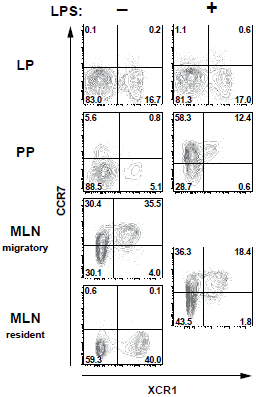
XCR1-expressing cells upregulate CCR7 after inflammation in Peyer’s patches and MLN. C57BL/6 mice were injected i. p. with LPS or with PBS for control, and the expression of CCR7 and XCR1 was compared 14 h later on DC (CD45^+^ Lin^-^ F4/80^-^ CD11c^+^ MHCII^+^ cells) in LP, PP, and MLN. Shown is one representative experiment out of three.

### XCR1^+^ migratory DC excel in cross-presentation of orally applied antigen

In order to test the ability of various intestinal DC populations to cross-present orally applied antigen, mice were fed with 25 mg of soluble ovalbumin (OVA), sacrificed 17 h later, and the various intestinal DC subsets isolated to high purity (> 98.5%). The DC subsets were then co-cultured at various ratios with OT I T cells to test their capacity to cross-present the OVA-derived peptide SIINFEKL to CD8^+^ T cells. When the percentage of proliferating OT I T cells was determined after 2.5 days of culture, MLN migratory XCR1^+^ CD103^+^ DC performed best in activating OT I T cells in 5 out of 5 experiments, while migratory XCR1^-^ DC, irrespective of their CD103 expression, were less effective (Fig. 7). Resident MLN DC also had a low stimulatory effect on OT I T cells (Fig. 7). Interestingly, when LP DC subsets were tested in the same experiments and thus directly compared with MLN DC, essentially no proliferation of OT I T cells was seen (Fig. 7). Since these LP DC were fully capable to activate OT I T cells when in vitro loaded with the peptide SIINFEKL, these results suggested that LP DC did not present sufficient antigen to activate OT I T cells. Taken together, in all conditions, in which substantial cross-presentation could be observed, migratory XCR1^+^ DC outperformed all other DC populations in the cross-presentation of orally applied antigen.

**Figure 7.**
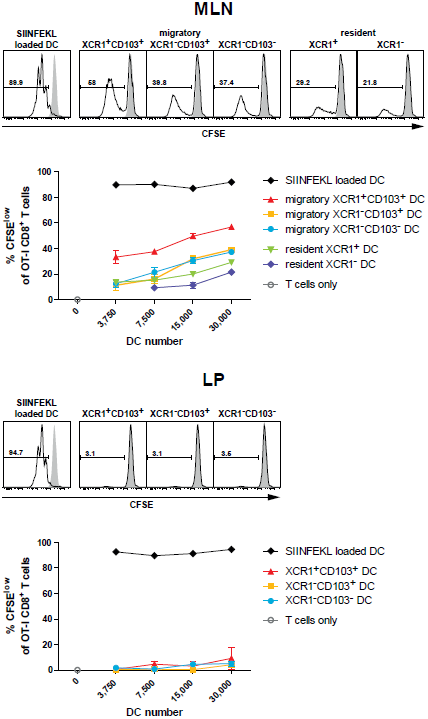
Intestinal XCR1^+^ migratory DC excel in cross-presentation of soluble antigen. C57BL/6 mice were fed with soluble 25 mg OVA, sacrificed 17 h later, and CD11c^+^ cells were enriched from MLN and LP by density gradient centrifugation and positive magnetic separation. The indicated DC subsets were flow-sorted according to their expression of XCR1 and CD103 to purity (>98.5%). Titrated numbers of the respective DC subsets from MLN or LP were then co-cultured with 1×10^5^ CFSE-labeled OT-I T cells, DC loaded with SIINFEKL peptide in vitro served as positive controls. Shown is the CFSE dilution profile of the OT-I T cells (CD90.1^+^ CD8^+^) after 2.5 d of co-culture with the respective DC subsets (open histograms), or OT I T cells alone (gray histograms). The results shown are representative of three experiments with LP and MLN, and two additional experiments with MLN only (mean ± SD).

## Discussion

We have previously extensively characterized splenic XCR1^+^ DC using a variety of approaches. All results obtained were compatible with the notion that the surface expression of the chemokine receptor XCR1 characterizes the Batf3-dependent lineage of cross-presenting DC (Bachem et al., 2012). In the present work we sought to determine whether this finding can be extended to the intestinal immune system.

Several levels of evidence indicate that XCR1^+^ DC also in the intestine are a separate lineage of DC with a consistent phenotype and a special ability to cross-present antigen. First of all, in animals deficient for the TF Batf3 only XCR1^+^ DC were consistently absent, while DC bearing other surface markers were preserved. In particular, XCR1^+^ CD103^+^ and XCR1^+^ CD103^-^ DC were absent, while XCR1^-^ CD103^+^ DC remained present, and the same was true for DC double-negative for CD103 and XCR1. Thus, in Batf3-deficient animals only the XCR1^+^ DC population failed to differentiate from precursors, while the development of XCR1^-^ DC remained intact.

Second, expression of XCR1 was in all instances closely correlated with the expression of CD8α and Clec9A/DNGR-1 (if detectable) on DC in all intestinal compartments and the draining MLN. Both CD8α and Clec9A/DNGR-1 have been identified in the past as integral parts of the gene expression programs of splenic cross-presenting DC and their putative correlates in the periphery (Crozat et al., 2010). This high correlation between expression of XCR1, Clec9A/DNGR-1, and CD8α in various lymphoid tissues strongly suggests a cooperation of these molecules in the specific function of cross-presenting DC. While the contribution of CD8α in this process remains unclear, XCR1 functions as the only receptor for XCL1 (Yoshida et al., 1998), a chemokine mainly released by innate cells (NK, NKT) and CD8^+^ T cells (Dorner et al., 2002; Dorner et al., 2004), and thus cells involved in the cytotoxic response, and more generally, in the type 1 immune defense. Clec9A/DNGR-1 has been described as a receptor of actin filaments expressed by necrotic cells, and is thought to contribute to the uptake of damaged cells into cross-presenting DC (Ahrens et al., 2012; Zhang et al., 2012). In spite of this close correlation between XCR1, CD8α, and Clec9A/DNGR-1 on (intestinal) DC, it has to be stated that only XCR1 is specifically expressed on conventional (intestinal) DC, while Clec9A/DNGR-1 is also present on plasmacytoid DC (pDC) (Caminschi et al., 2008; Sancho et al., 2008), and CD8α on pDC, T cells, NKT cells and other cells.

Third, our data show that XCR1^+^ DC excel in cross-presentation also in the intestinal immune system. Only very few cross-presentation experiments have been performed with intestinal DC in the past, in particular after application of antigen via the oral route. Chung et al. (Chung et al., 2005) reported that only CD11c^+^ DC isolated from the MLN were capable to cross-present orally applied OVA. At the same time, they somewhat surprisingly found that CD8^-^ CD11b^+^ DCs (i.e. XCR1^-^) rather than CD8^+^ DCs (i.e. largely XCR1^+^) cross-present intestinal Ag. When Jaensson et al. (Jaensson et al., 2008) re-investigated this issue, only CD103^+^ DC isolated from MLN after oral administration of OVA induced proliferation of OT I T cells in vitro. However, these authors did not further discriminate between CD103^+^ CD11b^-^ (“CD8α-like”, “Batf3-dependent”) and CD103^+^ CD11b^+^ (“Batf3-independent”) DC in their study, and therefore their results, from today’s perspective, did not offer the required DC subset resolution. In our work, we have separately tested LP and both migratory and resident MLN DC, and also subdivided these populations into XCR1^+^ and XCR1^-^ DC. DC originating from the LP usually did not give a signal in the cross-presentation assay; possibly the proportion of DC which have taken up antigen was too low, or the isolation procedure introduced a bias into the DC population. When analyzing MLN populations, the migratory DC clearly outperformed the resident DC, which still gave a low signal. Within the migratory DC population, the XCR1^+^ DC consistently performed best in the cross-presentation of orally applied soluble antigen, but a lower signal was also obtained with XCR1^-^ DC. These results were thus comparable to the data we have obtained earlier with splenic DC subsets. There, XCR1^+^ DC performed best, but not in a “unique fashion”, in the cross-presentation of soluble antigen (applied intravenously), but were “unique” in the cross-presentation of cell-bound antigen (Bachem et al., 2012). We did not have the opportunity to test the uptake and presentation of cell-bound antigen in our current work on intestinal DC. However, we are aware of the work of Cerovic et al. (submitted), who used for their studies a transgenic mouse in which only intestinal epithelial cells (IEC) express non-secretable antigen. In this system, Cerovic et al. found that only CD103^+^ CD11b^-^ DC (i.e. XCR1^+^ DC) migrating from the intestinal system, but not any other subset, were exclusively capable to cross-present this cell-associated antigen. Thus, combining the results of Cerovic et al. on cell-bound antigen with our own results on the presentation of orally applied antigen, we can clearly conclude that XCR1^+^ / CD103^+^ CD11b^-^ are the population of DC which excels in antigen cross-presentation in the intestinal immune system. This stated, one should bear in mind that these two designation systems define largely overlapping, but not fully congruent DC populations; essentially all CD103^+^ CD11b^-^ DC bear the XCR1 receptor, but XCR1 is also expressed on 10%-15% of other DC subdivided by the CD103/CD11b classification system (compare Fig. 1).

The specific expression of XCR1 allowed us to unequivocally determine the anatomical localization of cross-presenting DC in the intestinal system. In the small intestine, cells expressing XCR1^+^ were found in the lamina propria of the villi and also in epithelial crypts. In PP, XCR1^+^ DC mainly clustered in the interfollicular region, and were absent from the subepithelial dome. The latter results are congruent with the findings of Iwasaki et al. (Iwasaki and Kelsall, 2000), who, based on double-fluorescence studies using CD11c and CD8α as markers, described the presence of CD8α DC in the interfollicular region. Interestingly, XCR1^+^ signals were absent in the subepithelial dome of the PP, where CD11c^+^ CD11b^+^ cells can be found instead (Iwasaki and Kelsall, 2000). This anatomical separation in the PP indicates a division of labor of DC subsets in this lymphoid organ which at present is not fully understood. Finally, the distribution of XCR1^+^ DC in the draining MLN followed the pattern obtained with peripheral LN obtained earlier (Dorner et al., 2009), with signals present in T cell zones and apparently sinuses, where incoming (cellular) material in the draining lymph can be taken up by DC.

Our results on the phenotype, differentiation, function, and localization of DC demonstrate that XCR1^+^ DC are a homogenous population with a specific function in the intestinal immune system. With this understanding, we sought to determine, whether there are any surface molecules which would fully demarcate XCR1^-^ DC in a positive fashion. Interestingly, of the many markers tested, there were only two surface molecules which in all instances and in all tissues were not expressed on XCR1^+^ DC, namely CD11b and CD172a/SIRPα. When analyzing DC for the expression of these two integrins, it became apparent that all of XCR1^-^ DC express SIRPα, and some of them also CD11b (compare Fig. 2). Thus, only SIRPα showed a stringent anti-correlation with XCR1 on DC in the intestine, strongly suggesting that this surface molecule comprehensively characterizes the XCR1^-^ DC population. Support for this conclusion comes from our studies in the spleen, where the same constellation was found (Bachem et al., 2012), and our preliminary data indicate that this is also true in other body compartments (own unpublished results).

SIRPα, an Ig-superfamily transmembrane protein, is in the immune system abundantly expressed on macrophages, DC, and neutrophils; outside of the immune system it is present on neurons and weakly also on fibroblasts and endothelial cells (Matozaki et al., 2009; Nuvolone et al., 2013). Although the function of SIRPα is not fully understood, it has been implicated in the control of cell phagocytosis. Cells expressing CD47, the ligand for SIRPα, appear to be protected from engulfment by phagocytic cells (Matozaki et al., 2009). It is intriguing to note that Clec9A/ DNGR-1 and SIRPα, which on DC are never co-expressed, both regulate cell phagocytosis. This functional feature possibly contributes to the division of labor between the XCR1^+^ and SIRPα^+^ DC populations.

Future work will determine whether SIRPα^+^ DC are a homogenous population or whether they have to be further split up into functional subsets. For practical purposes it seems attractive now to classify DC in the immune system based on the expression of XCR1 and SIRPα, which greatly simplifies any phenotypical and functional analysis when compared with the DC classification systems currently in use. Figure 8 illustrates such an approach with DC populations from the various compartments of the intestine. Once separated into XCR1^+^ versus SIRPα^+^ positive DC, these subpopulations can be further analyzed for expression of other molecules which are being actively regulated in various tissue compartment, e.g. CD103 (Sathe et al., 2011; Zhan et al., 2011). It appears likely that this approach for the functional subdivision of DC can universally be applied throughout the immune system.

**Figure 8.**
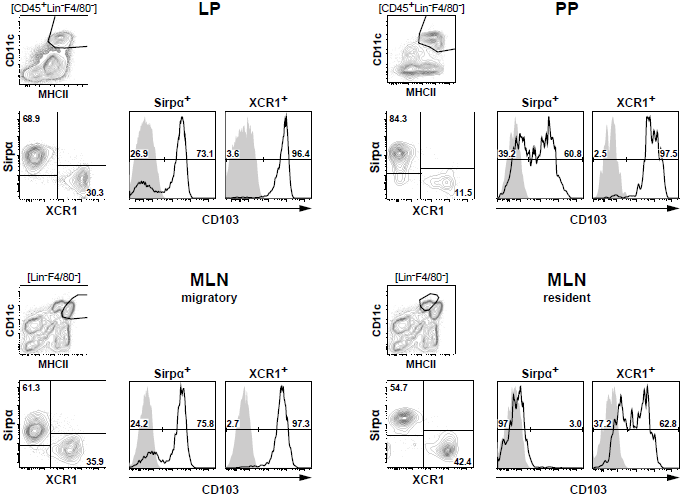
Classification of intestinal DC according to their expression of XCR1 and SIRPα. DC from the LP, PP, and MLN of C57BL/6 mice were enriched by digestion and density gradient centrifugation of the tissues, stained for XCR1 and SIRPα, DC from MLN were further separated into resident and migratory DC based on their MHCII expression levels. For analysis, the gates were set on live CD45^+^ Lin^-^ F4/80^-^ CD11c^+^ MHCII^+^ cells. Expression of CD103 is shown on XCR1^+^ versus SIRPα^+^ DC. The background staining was determined with homozygous B6.XCR1-lacZ^+/+^ mice lacking XCR1 (gray). The results shown are representative of two experiments.

## Author contributions

M.B. and S.Gü. designed and did experiments, analyzed and interpreted the data and contributed to the writing of the manuscript; A.B., E.H., A.M., A.J., A.H., V.H., H.W.M. and S.G. provided specialized expertise and/or assisted with some experiments; R.A.K. conceived the project, interpreted the data and wrote the manuscript.

## Acknowledgement

We are grateful to Maria Rescigno and her laboratory for introducing us to the methodology of cell isolation from intestinal tissues compartments. Batf3-deficient mice were kindly provided by H.-C. Probst, Mainz University, CX_3_CR1^GFP^ mice by M. Gunzer (Magdeburg), and Langerin^EGFP^ mice by P. Stoitzner (Innsbruck). Anti-murine Clec9A/DNGR 1 mAb were kindly provided by M. H. Lahoud and I. Caminschi (Melbourne) and C. Reis e Sousa (London). B16 cells secreting Flt3 ligand were a gift of S. Jung (Rehovot). We thank Reinhard Pabst, Hannover, for discussions on the positioning of XCR1^+^ DC in the intestinal immune tissues. This work was supported by the Fritz-Thyssen-Foundation, the Wilhelm Sander-Foundation, and the Deutsche Forschungsgemeinschaft (Kr 827/16-1 and 827/18-1).

## References

Ahrens, S., Zelenay, S., Sancho, D., Hanc, P., Kjaer, S., Feest, C., Fletcher, G., Durkin, C., Postigo, A., Skehel, M., Batista, F., Thompson, B., Way, M., Reis E Sousa, C., and Schulz, O. (2012). F-actin is an evolutionarily conserved damage-associated molecular pattern recognized by DNGR-1, a receptor for dead cells. Immunity 36, 635–645. doi:10.1016/j.immuni.2012.03.008

Annacker, O., Coombes, J.L., Malmstrom, V., Uhlig, H.H., Bourne, T., Johansson-Lindbom, B., Agace, W.W., Parker, C.M., and Powrie, F. (2005). Essential role for CD103 in the T cell-mediated regulation of experimental colitis. J. Exp. Med. 202, 1051–1061. doi:10.1084/ jem.20040662

Bachem, A., Güttler, S., Hartung, E., Ebstein, F., Schaefer, M., Tannert, A., Salama, A., Movassaghi, K., Opitz, C., Mages, H.W., Henn, V., Kloetzel, P.M., Gurka, S., and Kroczek, R.A. (2010). Superior antigen cross-presentation and XCR1 expression define human CD^-^ 11c^+^CD141^+^ cells as homologues of mouse CD8^+^ dendritic cells. J. Exp. Med. 207, 1273–1281. doi:10.1084/jem.20100348

Bachem, A., Hartung, E., Güttler, S., Mora, A., Zhou, X., Hegemann, A., Plantinga, M., Mazzini, E., Stoitzner, P., Gurka, S., Henn, V., Mages, H.W., and Kroczek, R.A. (2012). Expression of XCR1 characterizes the Batf3-dependent lineage of dendritic cells capable of antigen cross-presentation. Front. Immunol. 3, 214. doi:10.3389/fim-mu.2012.00214

Bar-On, L., Birnberg, T., Lewis, K.L., Edelson, B.T., Bruder, D., Hildner, K., Buer, J., Murphy, K.M., Reizis, B., and Jung, S. (2010). CX_3_ CR1^+^ CD8α^+^ dendritic cells are a steady-state population related to plasmacytoid dendritic cells. Proc. Natl. Acad. Sci. USA 107, 14745–14750. doi:10.1073/pnas.1001562107

Bevan, M.J. (2006). Cross-priming. Nat. Immunol. 7, 363–365. doi:10.1038/ni0406-363

Bogunovic, M., Mortha, A., Muller, P.A., and Merad, M. (2012). Mononuclear phagocyte diversity in the intestine. Immunol. Res. 54, 37–49. doi:10.1007/s12026-012-8323-5

Caminschi, I., Proietto, A.I., Ahmet, F., Kitsoulis, S., Shin, T.J., Lo, J.C., Rizzitelli, A., Wu, L., Vremec, D., Van Dommelen, S.L., Campbell, I.K., Maraskovsky, E., Braley, H., Davey, G.M., Mottram, P., Van, D.V., Jensen, K., Lew, A.M., Wright, M.D., Heath, W.R., Shortman, K., and Lahoud, M.H. (2008). The dendritic cell subtype-restricted C-type lectin Clec9A is a target for vaccine enhancement. Blood 112, 3264–3273. doi:10.1182/blood-2008-05-155176

Chung, Y., Chang, J.H., Kweon, M.N., Rennert, P.D., and Kang, C.Y. (2005). CD8α-11b^+^ dendritic cells but not CD8α^+^ dendritic cells mediate cross-tolerance toward intestinal antigens. Blood 106, 201–206

Crozat, K., Guiton, R., Contreras, V., Feuillet, V., Dutertre, C.A., Ventre, E., Vu Manh, T.P., Baranek, T., Storset, A.K., Marvel, J., Boudinot, P., Hosmalin, A., Schwartz-Cornil, I., and Dalod, M. (2010). The XC chemokine receptor 1 is a conserved selective marker of mammalian cells homologous to mouse CD8α^+^ dendritic cells. J. Exp. Med. 207, 1283–1292. doi:10.1084/jem.20100223

Crozat, K., Tamoutounour, S., Vu Manh, T.P., Fossum, E., Luche, H., Ardouin, L., Guilliams, M., Azukizawa, H., Bogen, B., Malissen, B., Henri, S., and Dalod, M. (2011). Cutting Edge: Expression of XCR1 defines mouse lymphoid-tissue resident and migratory dendritic cells of the CD8α^+^ type. J. Immunol. 187, 4411–4415. doi:10.4049/jim-munol.1101717

Den Haan, J.M., Lehar, S.M., and Bevan, M.J. (2000). CD8^+^ but not CD8^-^ dendritic cells cross-prime cytotoxic T cells in vivo. J. Exp. Med. 192, 1685–1696

Dorner, B.G., Dorner, M.B., Zhou, X., Opitz, C., Mora, A., Güttler, S., Hutloff, A., Mages, H.W., Ranke, K., Schaefer, M., Jack, R.S., Henn, V., and Kroczek, R.A. (2009). Selective expression of the chemokine receptor XCR1 on cross-presenting dendritic cells determines cooperation with CD8^+^ T cells. Immunity 31, 823–833. doi:10.1016/j.immuni.2009.08.027

Dorner, B.G., Scheffold, A., Rolph, M.S., Hüser, M.B., Kaufmann, S.H., Radbruch, A., Flesch, I.E., and Kroczek, R.A. (2002). MIP-1α, MIP-1β, RANTES, and ATAC/lymphotactin function together with IFN-γ as type 1 cytokines. Proc. Natl. Acad. Sci. USA 99, 6181–6186

Dorner, B.G., Smith, H.R., French, A.R., Kim, S., Poursine-Laurent, J., Beckman, D.L., Pingel, J.T., Kroczek, R.A., and Yokoyama, W.M. (2004). Coordinate expression of cytokines and chemokines by NK cells during murine cytomegalovirus infection. J. Immunol. 172, 3119–3131

Edelson, B.T., Kc, W., Juang, R., Kohyama, M., Benoit, L.A., Klekotka, P.A., Moon, C., Albring, J.C., Ise, W., Michael, D.G., Bhattacharya, D., Stappenbeck, T.S., Holtzman, M.J., Sung, S.S., Murphy, T.L., Hildner, K., and Murphy, K.M. (2010). Peripheral CD103^+^ dendritic cells form a unified subset developmentally related to CD8α^+^ conventional dendritic cells. J. Exp. Med. 207, 823–836. doi:10.1084/ jem.20091627

Hashimoto, D., Miller, J., and Merad, M. (2011). Dendritic cell and macrophage heterogeneity in vivo. Immunity 35, 323–335. doi:10.1016/j.immuni.2011.09.007

Hildner, K., Edelson, B.T., Purtha, W.E., Diamond, M., Matsushita, H., Kohyama, M., Calderon, B., Schraml, B.U., Unanue, E.R., Diamond, M.S., Schreiber, R.D., Murphy, T.L., and Murphy, K.M. (2008). Batf3 deficiency reveals a critical role for CD8α^+^ dendritic cells in cytotoxic T cell immunity. Science 322, 1097–1100. doi:10.1126/science.1164206

Iwasaki, A., and Kelsall, B.L. (2000). Localization of distinct Peyer’s patch dendritic cell subsets and their recruitment by chemokines macrophage inflammatory protein (MIP)-3α, MIP-3β, and secondary lymphoid organ chemokine. J. Exp. Med. 191, 1381–1394

Jaensson, E., Uronen-Hansson, H., Pabst, O., Eksteen, B., Tian, J., Coombes, J.L., Berg, P.L., Davidsson, T., Powrie, F., Johansson-Lindbom, B., and Agace, W.W. (2008). Small intestinal CD103^+^ dendritic cells display unique functional properties that are conserved between mice and humans. J. Exp. Med. 205, 2139–2149. doi:10.1084/jem.20080414

Jang, M.H., Sougawa, N., Tanaka, T., Hirata, T., Hiroi, T., Tohya, K., Guo, Z., Umemoto, E., Ebisuno, Y., Yang, B.G., Seoh, J.Y., Lipp, M., Kiyono, H., and Miyasaka, M. (2006). CCR7 is critically important for migration of dendritic cells in intestinal lamina propria to mesenteric lymph nodes. J. Immunol. 176, 803–810

Johansson-Lindbom, B., Svensson, M., Pabst, O., Palmqvist, C., Marquez, G., Förster, R., and Agace, W.W. (2005). Functional specialization of gut CD103^+^ dendritic cells in the regulation of tissue-selective T cell homing. J. Exp. Med. 202, 1063–1073. doi:10.1084/jem.20051100

Jung, S., Aliberti, J., Graemmel, P., Sunshine, M.J., Kreutzberg, G.W., Sher, A., and Littman, D.R. (2000). Analysis of fractalkine receptor CX_3_CR1 function by targeted deletion and green fluorescent protein reporter gene insertion. Mol. Cell. Biol. 20, 4106–4114

Kelsall, B.L., and Strober, W. (1996). Distinct populations of dendritic cells are present in the subepithelial dome and T cell regions of the murine Peyer’s patch. J. Exp. Med. 183, 237–247

Kissenpfennig, A., Ait-Yahia, S., Clair-Moninot, V., Stossel, H., Badell, E., Bordat, Y., Pooley, J.L., Lang, T., Prina, E., Coste, I., Gresser, O., Renno, T., Winter, N., Milon, G., Shortman, K., Romani, N., Lebecque, S., Malissen, B., Saeland, S., and Douillard, P. (2005). Disruption of the langerin/CD207 gene abolishes Birbeck granules without a marked loss of Langerhans cell function. Mol. Cell. Biol. 25, 88–99. doi:10.1128/MCB.25.1.88-99.2005

Mach, N., Gillessen, S., Wilson, S.B., Sheehan, C., Mihm, M., and Dranoff, G. (2000). Differences in dendritic cells stimulated in vivo by tumors engineered to secrete granulocyte-macrophage colony-stimulating factor or Flt3-ligand. Cancer. Res. 60, 3239–3246

Macpherson, G.G., Jenkins, C.D., Stein, M.J., and Edwards, C. (1995). Endotoxin-mediated dendritic cell release from the intestine. Characterization of released dendritic cells and TNF dependence. J. Immunol. 154, 1317–1322

Maraskovsky, E., Brasel, K., Teepe, M., Roux, E.R., Lyman, S.D., Shortman, K., and Mckenna, H.J. (1996). Dramatic increase in the numbers of functionally mature dendritic cells in Flt3 ligand-treated mice: multiple dendritic cell subpopulations identified. J. Exp. Med. 184, 1953–1962

Matozaki, T., Murata, Y., Okazawa, H., and Ohnishi, H. (2009). Functions and molecular mechanisms of the CD47-SIRPα signalling pathway. Trends Cell. Biol. 19, 72–80. doi:10.1016/j.tcb.2008.12.001

Milling, S., Yrlid, U., Cerovic, V., and Macpherson, G. (2010). Subsets of migrating intestinal dendritic cells. Immunol. Rev. 234, 259–267. doi:10.1111/j.0105-2896.2009.00866.x

Mowat, A.M., and Bain, C.C. (2011). Mucosal macrophages in intestinal homeostasis and inflammation. J. Innate Immun. 3, 550–564. doi:10.1159/000329099

Nuvolone, M., Kana, V., Hutter, G., Sakata, D., Mortin-Toth, S.M., Russo, G., Danska, J.S., and Aguzzi, A. (2013). SIRPα polymorphisms, but not the prion protein, control phagocytosis of apoptotic cells. J. Exp. Med. 210, 2539–2552. doi:10.1084/jem.20131274

Ohl, L., Mohaupt, M., Czeloth, N., Hintzen, G., Kiafard, Z., Zwirner, J., Blankenstein, T., Henning, G., and Förster, R. (2004). CCR7 governs skin dendritic cell migration under inflammatory and stea-dy-state conditions. Immunity 21, 279–288. doi:10.1016/j.immu-ni.2004.06.014

Pabst, O., and Bernhardt, G. (2010). The puzzle of intestinal lamina propria dendritic cells and macrophages. Eur. J. Immunol. 40, 2107–2111. doi:10.1002/eji.201040557

Pooley, J.L., Heath, W.R., and Shortman, K. (2001). Cutting edge: intravenous soluble antigen is presented to CD4 T cells by CD8^-^ dendritic cells, but cross-presented to CD8 T cells by CD8^+^ dendritic cells. J. Immunol. 166, 5327–5330

Rescigno, M. (2010). Intestinal dendritic cells. Adv. Immunol. 107, 109–138. doi:10.1016/B978-0-12-381300-8.00004-6

Sancho, D., Mourao-Sa, D., Joffre, O.P., Schulz, O., Rogers, N.C., Pennington, D.J., Carlyle, J.R., and Reis E Sousa, C. (2008). Tumor therapy in mice via antigen targeting to a novel, DC-restricted C-type lectin. J. Clin. Invest. 118, 2098–2110. doi:10.1172/JCI34584

Sanes, J.R., Rubenstein, J.L., and Nicolas, J.F. (1986). Use of a recombinant retrovirus to study post-implantation cell lineage in mouse embryos. EMBO J. 5, 3133–3142

Sathe, P., Pooley, J., Vremec, D., Mintern, J., Jin, J.O., Wu, L., Kwak, J.Y., Villadangos, J.A., and Shortman, K. (2011). The acquisition of antigen cross-presentation function by newly formed dendritic cells. J Immunol. doi:10.4049/jimmunol.1002683

Schulz, O., Jaensson, E., Persson, E.K., Liu, X., Worbs, T., Agace, W.W., and Pabst, O. (2009). Intestinal CD103^+^, but not CX3CR1^+^, antigen sampling cells migrate in lymph and serve classical dendritic cell functions. J. Exp. Med. 206, 3101–3114. doi:10.1084/jem.20091925

Shen, L., and Rock, K.L. (2006). Priming of T cells by exogenous antigen cross-presented on MHC class I molecules. Curr. Opin. Immunol. 18, 85-91. doi:10.1016/j.coi.2005.11.003

Turnbull, E.L., Yrlid, U., Jenkins, C.D., and Macpherson, G.G. (2005). Intestinal dendritic cell subsets: differential effects of systemic TLR4 stimulation on migratory fate and activation in vivo. J. Immunol. 174, 1374–1384

Villadangos, J.A., Heath, W.R., and Carbone, F.R. (2007). Outside looking in: the inner workings of the cross-presentation pathway within dendritic cells. Trends Immunol. 28, 45–47. doi:10.1016/j.it.2006.12.008

Worbs, T., Bode, U., Yan, S., Hoffmann, M.W., Hintzen, G., Bernhardt, G., Förster, R., and Pabst, O. (2006). Oral tolerance originates in the intestinal immune system and relies on antigen carriage by dendritic cells. J. Exp. Med. 203, 519–527. doi:10.1084/jem.20052016

Yoshida, T., Imai, T., Kakizaki, M., Nishimura, M., Takagi, S., and Yoshie, O. (1998). Identification of single C motif-1/lymphotactin receptor XCR1. J. Biol. Chem. 273, 16551–16554

Zhan, Y., Carrington, E.M., Van Nieuwenhuijze, A., Bedoui, S., Seah, S., Xu, Y., Wang, N., Mintern, J.D., Villadangos, J.A., Wicks, I.P., and Lew, A.M. (2011). GM-CSF increases cross presentation and CD103 expression by mouse CD8^+^ spleen dendritic cells. Eur J Immunol. doi:10.1002/eji.201141540

Zhang, J.G., Czabotar, P.E., Policheni, A.N., Caminschi, I., San Wan, S., Kitsoulis, S., Tullett, K.M., Robin, A.Y., Brammananth, R., Van Delft, M.F., Lu, J., O‘reilly, L.A., Josefsson, E.C., Kile, B.T., Chin, W.J., Mintern, J.D., Olshina, M.A., Wong, W., Baum, J., Wright, M.D., Huang, D.C., Mohandas, N., Coppel, R.L., Colman, P.M., Nicola, N.A., Shortman, K., and Lahoud, M.H. (2012). The dendritic cell receptor Clec9A binds damaged cells via exposed actin filaments. Immunity 36, 646–657. doi:10.1016/j.immuni.2012.03.009

